# Rethinking the functions of peacock’s display and lek organisation in native populations of Indian Peafowl *Pavo cristatus*

**DOI:** 10.1101/2022.09.21.508866

**Authors:** Dhanashree Ashok Paranjpe, Vedanti Rajiv Mahimkar, Priyanka Dange

## Abstract

Males of Indian Peafowl are known for their extravagant courtship display used to attract mates. The displays are seen in absence of potential mates as well. This study investigates various contexts in which display behaviours are shown by Indian peafowl. Upto 70% of displays were observed in absence of potential mates. High frequency of displays in presence of other males and longer display bouts during the month of May when mating rarely happens, indicated that display behaviours might have a role to play in male-male competition or territory defence in addition to the female mate choice. The males establish and maintain their courtship display territories throughout the breeding season. Earlier studies assume that the choice of location does not depend on any resources as the species has a lek mating system. How these display sites are chosen is still a question. In this study, we investigated factors that might be important for the selection of display sites in a free-ranging Indian Peafowl population. It was observed that display sites were more concentrated within a radius of 300 meters of the food provisioning site and/ or a water resource while the number of display sites decreased considerably beyond the 300 m radius of food and water resources. Overall, the selection of display sites is non-random and highlights the importance of resources in the choice of display territories. The spatial organisation of leks in our study indicate that the mating system in Indian peafowl may be resource-based lek.

## Introduction

In many species, displays or behaviours involving secondary sexual characters may serve different functions such as a signal of aggression (Rowher 1982, Loyau et al., 2005), for the establishment and/or maintaining a territory (Wiens and Tuschhoff, 2020), male-male competition (Rowher 1982; Mateos, 1998; Loyau et al., 2005; Wiens and Tuschhoff, 2020) and as a dynamic character which plays an important role in female mate choice (Darwin 1871; Petrie et al.,1991; Candolin, 2003; Wiens and Tuschhoff, 2020).

Males of Indian peafowl *Pavo cristatus* are well known for extravagant displays involving mainly their colourful upper tail coverts which are used by females to choose a mate (Petrie et al., 1991; Petrie and Halliday 1994; Loyau et al. 2005, 2007; Dakin and Montgomerie, 2009). However, in this species display behaviours have been observed in absence of females as well (Yasmin and Yahya, 1996; Dakin and Montgomerie, 2009). The proportion of time spent in display in absence of females was estimated to be 24% of the active display time spent by a male on its display territory (Dakin and Montgomerie, 2009). Engaging in complex display behaviours for longer duration is thought to be energetically costly, therefore, a substantial proportion of active time spent in display in absence of females (for whom the behaviour is supposedly intended) warrants some explanation. According to some previous studies, territorial peacocks use their train as a threat in aggressive interactions to chase floating males without physical contact (Petrie et al., 1991, Loyau, 2005). Territorial birds might benefit from displaying their status or territory ownership to avoid escalation and related costs of energy expenditure as well as the risk of injury (Rowher, 1982). In red-necked pheasant *Phasianus colchicus* male traits such as tail and spur length, and wattle display duration was used in both intra- and inter-sexual selection (Reviewed by Mateos, 1998). The presence of other individuals of the same or opposite sex around the display territory may influence the display behaviours and/ or display might be used in the context(s) other than female choice. The proportion of active time spent in display in the presence or absence of other individuals might give clues about additional functions of display. The present study estimated the percentage of displays in the presence/ absence of other individuals of the same/ different sex in native populations of Indian peafowl.

Indian peafowl has been reported to have a lek mating system (Petrie et al.,1991, Yasmin and Yahya 1996; Loyau et al., 2005, 2007). Bradbury (1981) defined the following characteristics of a classic lek: i) There is no male parental care: Males contribute nothing to the next generation except gametes. ii) There is an arena or lek to which females come and on which most of the mating occurs. An arena is a site on which several males aggregate and that does not fill the habitat normally used by the species for other activities such as feeding, roosting, etc. iii) The display sites of males contain no significant resources required by females except the males themselves. This stipulation includes food, water, roosts, nest sites, egg deposition sites, etc. iv) The female has an opportunity to select a mate once she visits the arena.

Moreover, it is assumed that in a lekking species, the female choice of which male(s) to mate with is based solely on characteristics of the male and not related to direct fitness gains to females such as access to resources (Reynolds and Gross 1990; Hogland and Alatalo, 1995). Bradbury (1981) defined a continuum of male dispersion patterns during breeding season: from males defending resources crucial to females (resources defence polygyny), through a partially clustered ‘exploded or quasi lek’ to the highly clustered structure of classical lek. In exploded leks, males are often barely within visual or hearing distance of each other and the clustering is only apparent with careful mapping of display territories. Additionally, neither male attractiveness nor the spatial distribution of display territories on exploded leks depends on resources or males establish display territories in the areas that maximise female encounters, in accordance with the hotspot model of lek evolution (Bradbury and Gibson, 1983; Jiguet et al., 2002). On the other hand, in a resource-based lek, the spatial distribution of male display territories typically overlaps with resources crucial for females (food, nesting sites, etc.) but the male attractiveness is not related to the resources defended (Jiguet et al., 2002). For a resource-based lek, one would expect that the distribution of resources in the habitat will influence the choice of display territory during the breeding season. Under such a scenario, the spatial distribution of display sites will not be uniform or random, rather the display territories would be clustered around resources such as food and water. The possibility that resource distribution might influence lek organization in Indian peafowl was explored in a previous study by Loyau et al., 2007 to some extent. However, quantitative measurements of display territory size, lek size, and spatial distribution of display sites on a lek have not been done for populations of Indian peafowl in their native habitat. Spatial distribution of important resources and display territories would inform us about the lek organization of Indian peafowl in their native habitat.

The objectives of our study were:

1. To study if the choice of display territories by Indian peafowl males during breeding season is influenced by proximity to important resources such as food, water and/or female movements?

2. To determine the degree of clustering of male display territories quantitatively based on male territory size, the number of males per cluster and the distance between the clusters.

3. To understand the influence of other individuals (males, females, sub-adult males) on the display bouts of breeding males.

4. To understand the role of factors such as time of day and progression of the breeding season on the display bout frequency and duration.

## Materials and methods

Display bouts of males were recorded at various field sites across Rajasthan and Maharashtra from May 2016-Sept 2021 using opportunistic sampling. Details of field sites can be found in Table 1.

**Table 1.**
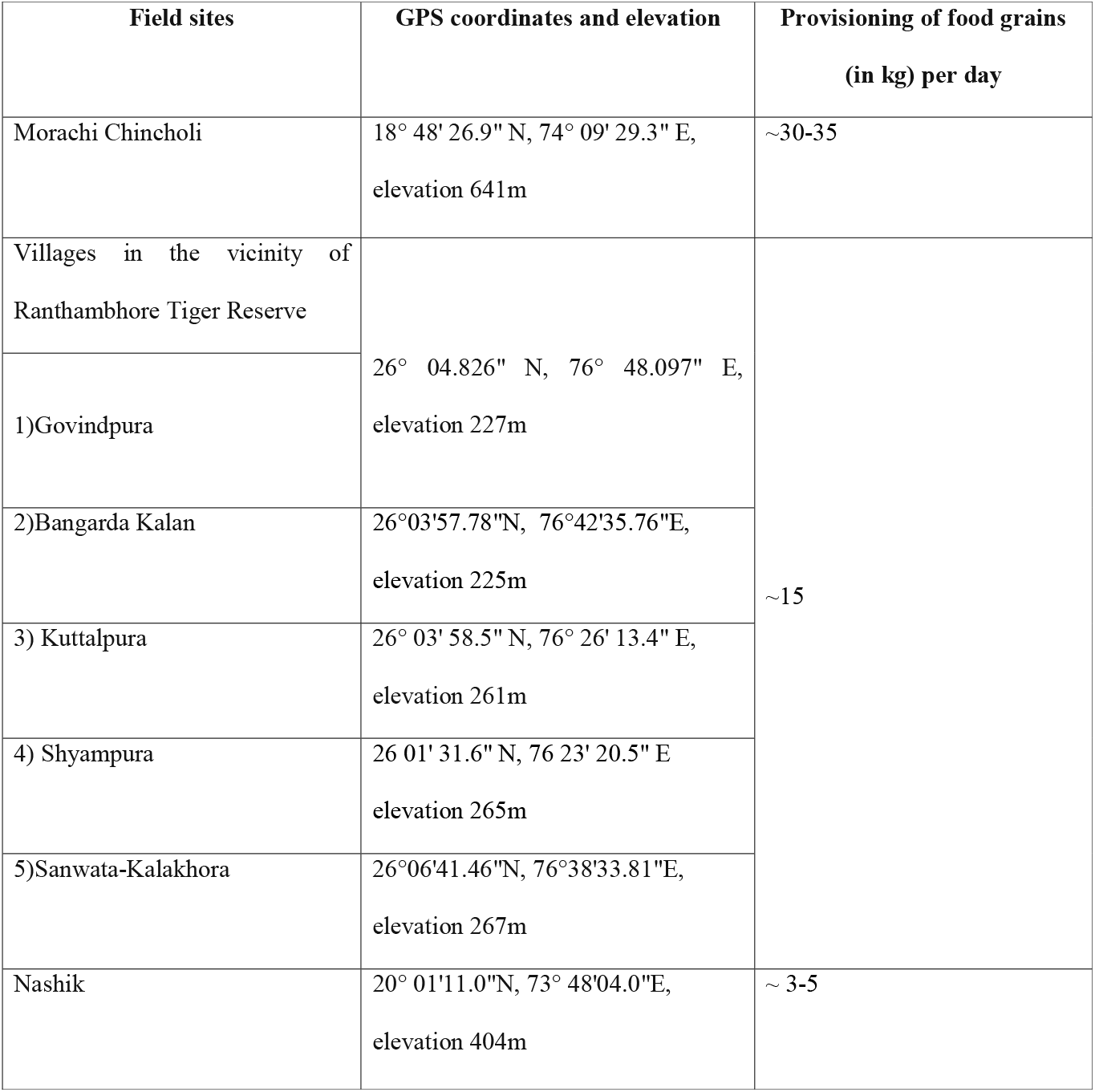
Brief description of field sites

The location of the display, date, time of the day, and start and ending time of each display bout was recorded. A total of 481 display bouts were recorded for 206 males during the study. Additionally, a record was kept if any other adult male(s), female(s) or sub-adult male(s) were seen within 50m distance of the focal displaying male (visibility at study sites during the breeding season is not more than 50-100m due to topography, growing crops, human habitations, etc.). If a female approached within 5m or closer of the displaying male it was recorded as ‘female visitation’ to the display rather than just the presence of a female around the focal male (anywhere within a 50m radius of the displaying male). To understand if the presence of other individuals changed the frequency and/or duration of focal male display bouts, we monitored males (N=28) for a minimum of 30 min or more till they kept displaying in that observation session and calculated the display frequency (display bouts per hour) as well as % display time (proportion of total observation time that they spent in display behaviour).

Physical characteristics and spatial distribution of display territories were studied only at one field site: Morachi Chincholi, Maharashtra (18° 48’ 26.9” N, 74° 09’ 29.3” E, elevation 641m) from 2016 to 2019 and 2021 (field data could not be collected during the year 2020 due to restrictions related to COVID-19 pandemic). Morachi Chincholi is a village in the state of Maharashtra, India that hosts a wild population of approximately 500-600 Indian peafowl in its agricultural landscape in close association with human habitation (based on unpublished data of population surveys done by authors in 2016, 2017, 2019). According to Paranjpe and Dange, 2020, approximately 30 to 35kg of food grains are offered at Morachi Chincholi per day at specific places in the village throughout the year which are referred here as ‘food provision sites’. The GPS locations of food provision sites were marked. Additionally, peafowl also has free access to agricultural lands surrounding the village and well-clumped dwellings that constitute the village. GPS locations of water resources such as wells, irrigation canals, farm ponds, percolation ponds, water troughs, and hand pumps were also marked and mapped using QGIS. GPS locations of individual or groups of peafowl were marked throughout the village during non-breeding season in 2020 (January, February, September and October).

The breeding season of Indian peafowl starts in Morachi Chincholi around mid-April and lasts till mid-October. The field site (∼ 11Km^2^ area of the village and its surroundings) was scanned for displaying males starting April through October in the years 2016 to 2019 and 2021. As a male was located displaying, GPS coordinates of the location were recorded in each year of data collection. Even if we saw a male displaying at the same location more than once, the location was marked only once during one breeding season. Some locations were used by males for display year after year. A separate record was kept of such repeated locations of the display.

### Analysis

Multiple regression analysis was used to check if the presence of other individuals influenced display frequency (display bouts per hour) and % display time of the focal males. The number of adult males, females, and sub-adult males present in the vicinity of focal displaying male as well as the number of female visitations were used as regressors, while display frequency and % display time were used as dependent variables, in separate analyses. Univariate tests of significance were used to check which of the regressors had a significant effect on display frequency and % display time. Analysis of Variance (ANOVA) was carried out to determine if field sites and progression of the breeding season (month of the year), both categorical variables, had an effect on display bout frequency and % display time.

The locations of display territories, food provisioning sites and sources of water marked using GPS were put on the map of the field site using QGIS software. Whether the same male uses the same location for display year after year cannot be determined since no individual identification tags, colours, or rings were used for tracking during the study. First, we set out to analyse if the display territories were uniformly distributed across the field site or formed distinct clusters? To answer this question, all display territories recorded across years were put on the map. Even though some of the display sites may not be used every year and few new sites might be added, many of the display locations across Morachi Chincholi remained more or less the same across the years. Hence, we pooled the location data for all years to know the spatial distribution of display territories at Morachi Chincholi. We tried to draw cluster polygons manually using the polygon tool in QGIS based on a visual estimate of which display territories formed a cluster. Seven distinct clusters of display territories were identified visually. For confirming these cluster assignments, we did cluster analysis in STATISTICA™ software (v.14.0.0.15, TIBCO Software Inc.) using the latitude and longitude coordinates of each display territory. For calculating initial cluster centres, observations were chosen by the software to maximize initial between-cluster distances. Spatial coordinates of the centroid (cluster centre) for each cluster polygon were extracted from the software. Euclidean distance between each display territory from the nearest food provision site within the cluster was also calculated in meters using QGIS.

To calculate the number of display sites in close vicinity of food provision sites, or water resources (irrigation canal, well, farm pond, etc.) buffer areas with 300m and 500m radii were drawn in the QGIS software keeping food provision sites as the centre. These buffer zones were overlaid on a map containing all display sites at Morachi Chincoli. Then no. of display sites within 0-300m zone, 300-500m zone and >500m zone from the feeding site were noted.

## Results

### Factors influencing Indian peafowl display

The males established display territory at the start of the breeding season and defended it throughout the breeding season. Males displayed at the defended location in the presence and/ or absence of females as well as other males. Out of 481 display bouts recorded during the study across Morachi Chincholi, Nashik and Rajasthan for 39.71% of bouts (N= 191), no other individual was observed around the displaying male. During 30.77% of bouts (N= 138), at least one adult male was present within 100m of the displaying male. 35.76% of displays (N= 172) were observed in presence of at least one female within a 50m distance of the displaying male. Female “visiting” a displaying male (defined as a female coming within 5m or closer of the displaying male) was observed in the case of 20.37% (N=98) of bouts. Males even displayed in presence of sub-adult males (17.67% of display bouts i.e., N= 85). Sometimes (5.2% of display bouts i.e., N=25) even sub-adult males were seen displaying even if they did not have fully developed train feathers. However, sub-adult males were not seen defending any display territory.

Displays by sub-adult males were often for a short duration (maximum 2-3 min.) and seen at random places including food provision sites, display territory of adult males, near female movement routes, etc. Display bout duration of adult males was different at different stages of the breeding season. According to the Median test, the duration of display bouts was significantly different according to the month of the year (Chi square= 34.62, d.f.= 9, *p*= 0.0001). Sporadic displays were observed starting in March, but these displays did not last longer than 2-3 minutes. Most of them lasted for less than a minute. Display bouts were longer in the months of May, June, and July, with a peak in the month of May across all field sites. Two-tailed multiple comparisons using Kruskal-Wallis test revealed that bout duration was significantly more in May compared to April (*p*= 0.01) and August (*p*= 0.000002), while bout duration in September was significantly more than that in August (*p*= 0.04). The duration of display bouts shortened in August when females generally start to lay eggs. September brings rains again in certain parts of India due to returning north-east monsoon. An increase in the display bout duration was observed in September across all field sites after which it tapered off till November. Around this time, males shed their feathers and the non-breeding season starts.

The display bout frequency (or display frequency) for males was calculated as the number of display bouts per hour based on at least 30 min. of observations of focal males at their display site (N=28). The display frequency varied from 2.12-12.33 bouts/ hour, with an average display frequency of 5.43 bouts/hr. The display frequency was significantly influenced by the progression of the breeding season (Univariate test of significance, *F*= 5.05, *d*.*f*.= 4, *p*= 0.005). Multiple comparisons using an unequal N HSD test revealed that display frequency in the month of August was significantly higher compared to that in the months of May (*p*= 0.008), June (*p*=0.007) and September (*p*=0.009).

The presence of other individuals can potentially influence how frequently and/ or how long males display. Multiple regression analysis using display frequency as the dependent variable (Whole model multiple R^2^= 0.44, *p*= 0.0098) showed that the number of other males present in the vicinity of focal displaying male significantly increased the display frequency (Univariate tests of significance; *p*= 0.0062), while female visitation significantly decreased the display frequency of focal male (Univariate tests of significance; *p*= 0.0047). The presence of females in the vicinity of displaying male significantly increases the percentage of time they spend in displays (Whole model multiple R^2^= 0.39, *p*= 0.0095; Univariate tests of significance; *p*= 0.0022).

During behavioural observations of displaying males, it was noted that certain elements of the dynamic display were observed only in presence of females. For E.g., males perform the train rattle movement mainly in presence of females by rapidly vibrating the rectrices that support their train feathers in the upright position, such that those feathers make a noise audible from several metres away. This was also called shivering by Petrie et al. (1992) and Takahashi et al. (2008) (Dakin and Montgomerie, 2009). Males also bend slightly forward while rattling their train and turn less when a female is very close by. In absence of female wing-shaking displays i.e., rapid up and down movements of orange secondary feathers while the train is up happens more often (Dakin and Montgomerie 2009, Also supplementary video S1). This wing-shaking element is rarely followed by train rattle if females are not visiting the display. Males also turn more in all directions when females are not in the vicinity.

We have identified calls of Indian peafowl that are unique to the breeding season (Mahimkar and Paranjpe, manuscript in preparation). One of the calls (Honk-korr-ke or Korr-ke) given by peacocks is a display advertisement call and is given generally either at the beginning or end of the display bout. There is a distinct call given in presence of another adult or sub-adult male where the territory-holding male runs towards the potential intruder with rapid neck movements giving a Honk-Kaan or pe-kaan call simultaneously. This call can be considered a territory defence call. In presence of females, a hoot call (considered a copulation call) is given followed by a quick dash towards the female in an attempt to mate. A hoot call can also be heard in absence of females. This ‘deceptive copulation’ call has been reported earlier by Dakin and Montgomerie (2014). It can potentially attract females to the displaying male. Whether a female is visiting or absent, the hoot call is always given while the male is actively displaying. Thus, some of the calls can be considered as additional features of the dynamic display along with the complex sequence of movements by the displaying male.

### Physical characters and spatial distribution of display territories

The data for physical characters and spatial distribution of display territories were collected mostly in Morachi Chincholi, Maharashtra. The general habitat at Morachi Chincholi is open grassy scrubland with few patches of plantation and occasional tall trees or cultivated land. A clearing on the ground due to repeated trampling, impressions of peafowl feet in the soil, occasionally shed/ broken feathers and faecal matter or white marks formed by excretion of uric acid confirm the presence of a display site even in absence of a guarding male, although the location was marked as a display site only if a male was seen actually displaying at the location. Typically, a medium to big-sized tree, thickets, mud/ stone wall or solid structure was noted immediately behind/ adjacent to where peacocks displayed. Out of 150 display sites for which physical features were clearly noted down, 129 (86%) sites had a moderate to big tree or thicket as a feature of the display site, 43 sites (28.67%) had a built structure such as wall, house, well in the immediate background. 48 out of 150 display sites (32%) had a water source such as an irrigation canal, water spout, well, or water trough while 83 sites (55.33%) had a food source on-site in the form of agricultural produce or grains put out for the peafowl (food provision sites). Thus, the presence of moderate to big tree or thickets and the proximity of food and water source seem to be important physical features of the display site.

A total of 187 display sites were located between 2016 -2021 in the ∼ 11 Km^2^ area of Morachi Chincholi and mapped using QGIS. Seven distinct clusters of display sites were observed within this area (Fig. 1). The k-means cluster analysis confirmed that within-cluster Euclidean distances between display sites were significantly shorter than distances across clusters using both latitude and longitude as variables. This analysis confirmed that the visual assignment of display sites to seven distinct clusters was correct. The average distance between the clusters was 1637.49 ± 94.84m (Average ± SE).

**Figure 1:**
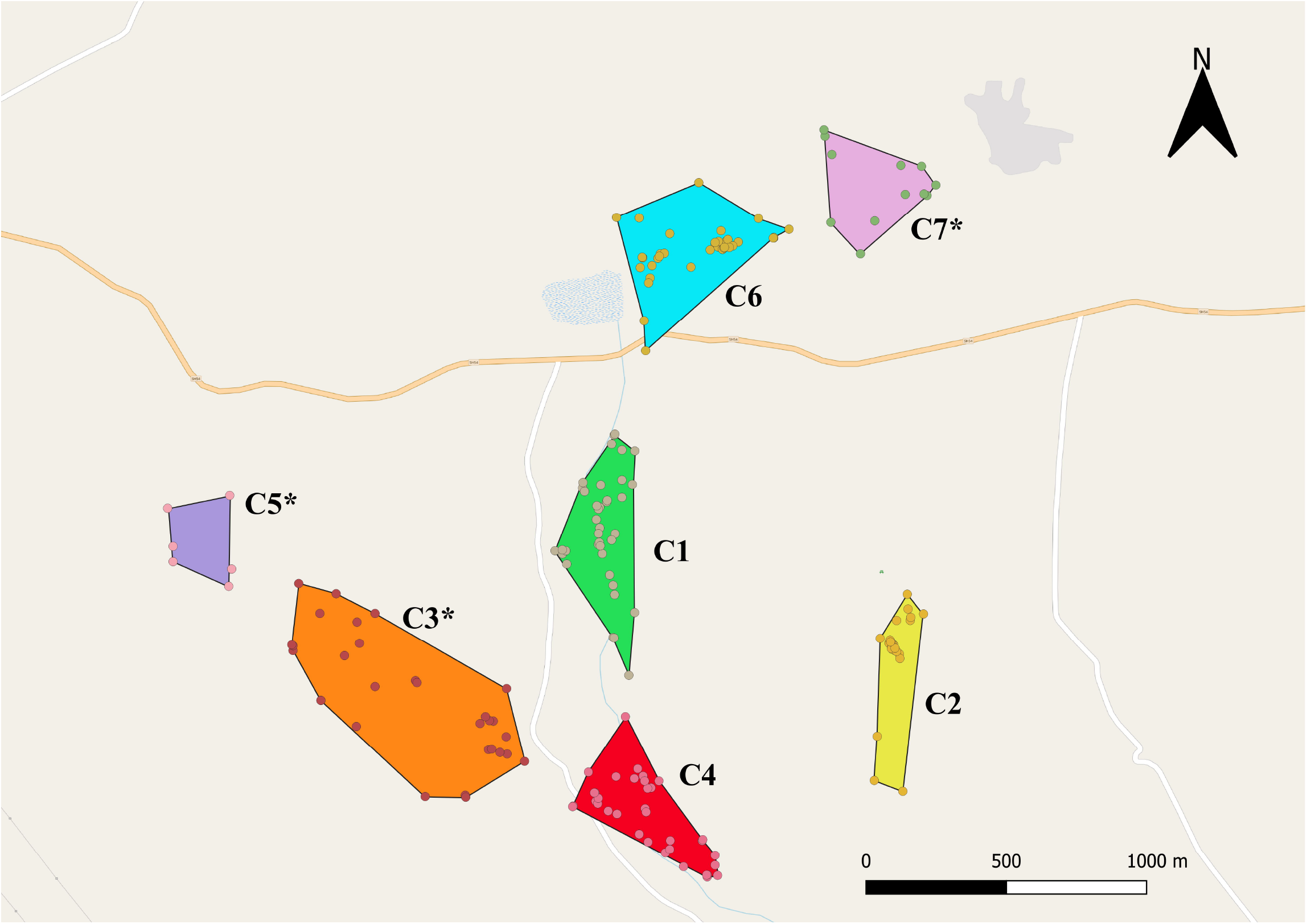
Display site clusters of Indian peafowl at Morachi Chincholi, Maharashtra. Seven clusters of display sites (C1 to C7) were spread over ∼11Km area. Clusters with asterisk (*) indicate the clusters which had no regular food provisioning site.

The number of display sites in clusters did not increase with the increase in the total area of the cluster (Table 2), however, the number of display sites showed a significant positive correlation with the number of water resources present within a cluster (Fig. 2). An irrigation canal runs through the field site starting from a percolation pond in the village (Fig. 3). Out of 187 display sites mapped during the study, 118 (63.1%) display sites were found within 500m zone on both sides of the irrigation canal. The remaining 69 display sites were within a 500m zone of at least one water resource other than an irrigation canal such as a well, hand pump, or farm pond (Fig 3).

**Table 2:**
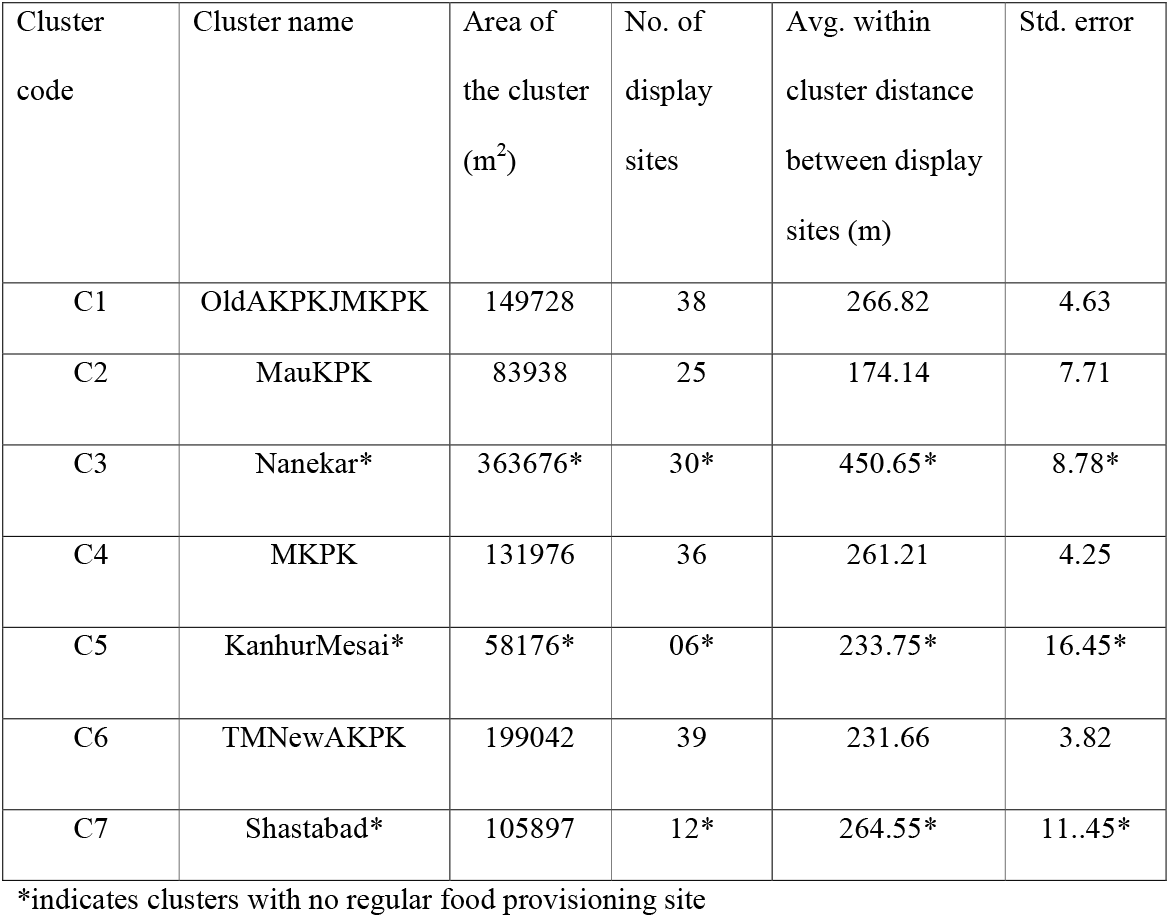
Summary of display site clusters in Morachi Chincholi.

**Fig. 2:**
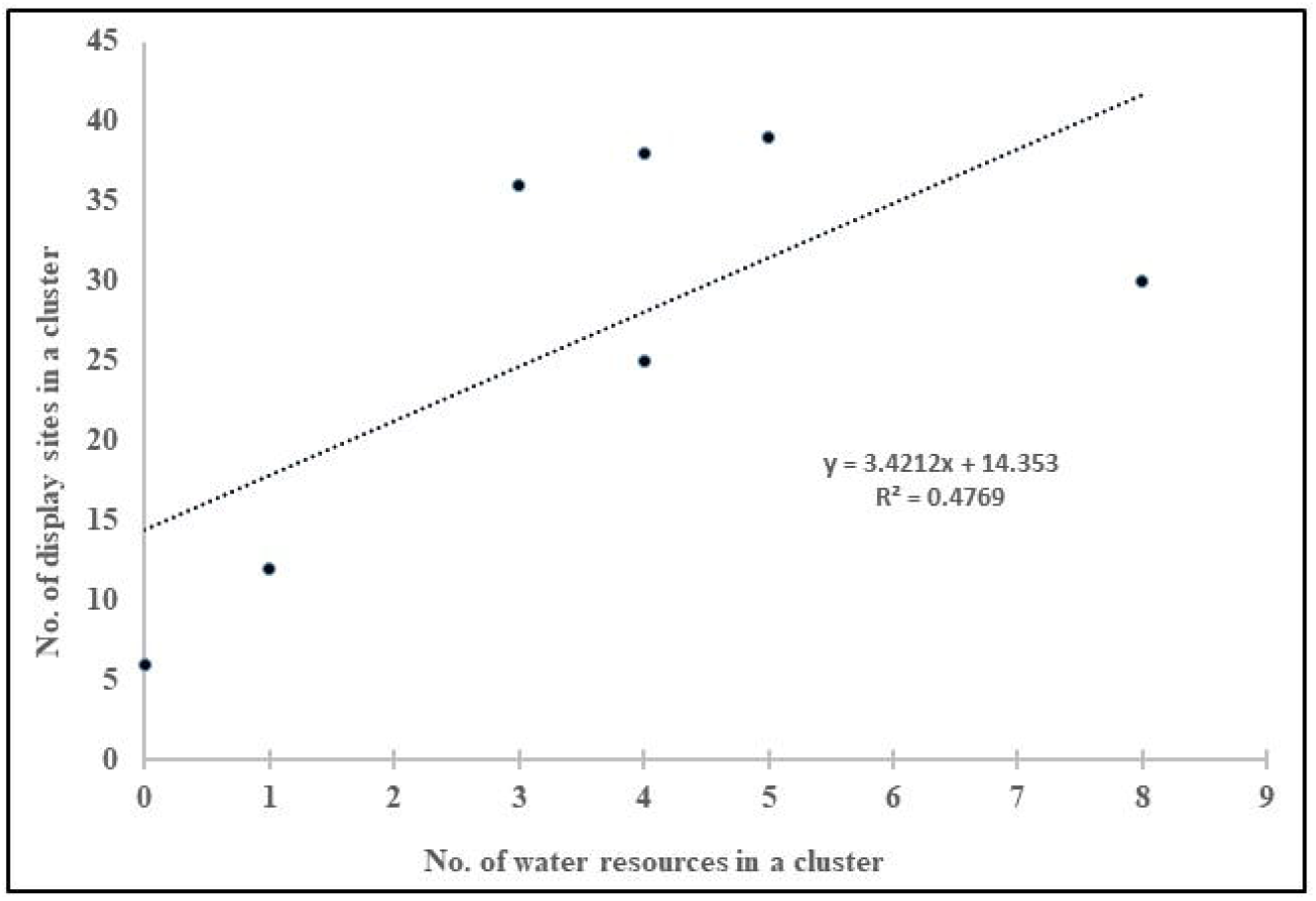
Correlation between no. of display sites with no. of water sources within each cluster

**Fig. 3:**
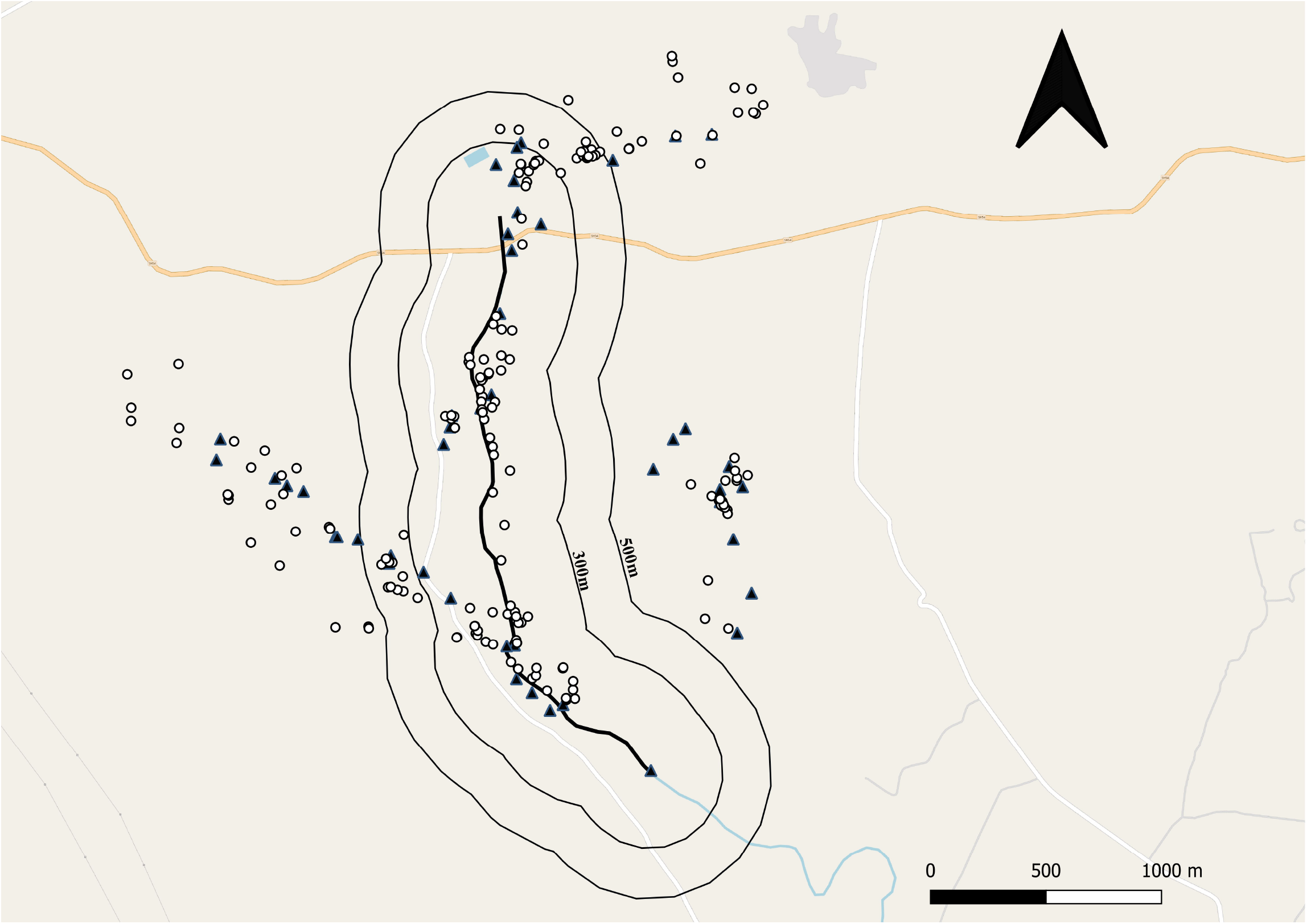
Distribution of display sites around the irrigation canal at Morachi Chincholi. The thick black line indicates the irrigation canal running through the field site. 300m and 500m zones on both sides around the canal are demarcated by contour lines. 63.1% of display sites (open circles) across years were found within 500m zone from the irrigation canal. Water resources other than irrigation canal are indicated by filled triangles.

Four clusters (C1, C2, C4, C6) had at least one regular food provisioning site within the cluster while three clusters (C3, C5 and C7) had no regular food provisioning site. The clusters that did not contain regular food provision sites had plenty of water sources such as wells, irrigation canal, hand pumps, and farm ponds just like other clusters. Kanhur mesai cluster (C5) had no water resources and no food provision sites within the cluster. Within each cluster, most of the display sites were located within 300-500m distance of the food provision sites (Table 3).

**Table 3:** Distance of individual display sites from a permanent food provision site within a cluster where grains are provided to the birds throughout the year.

| Cluster | Feeding site | No. of display sites within 0-300m of food provision site | No. of display sites within 301-500m of food provision site | No. of display sites within >500m of food provision site | Total no. of display sites in the cluster |
| --- | --- | --- | --- | --- | --- |
| C1 | F8 | 26 | 9 | 3 | 38 |
| C2 | F3 | 22 | 2 | 1 | 25 |
| C4 | F5 | 25 | 11 | 0 | 36 |
| C6 | F9 | 29 | 10 | 0 | 39 |

The area of individual display territories established by males ranged from 5.47m^2^ – 179.2m^2^. The average area of display territory in Morachi Chincholi was 33.02 ± 3.57 m^2^ (mean ± SE, N= 82). There was a weak negative correlation between the area of the display site and its distance from the nearest food provision site (N= 82, Correlation coefficient = -0.16, R^2^ = 0.0327).

## Discussion

The role of extravagant display of peacocks in attracting females and female mate choice has been the focus of many studies (Petrie et al, 1991; Yasmin and Yahya 1996; Loyau et al. 2005, 2007). However, our results showed that up to 70% of display bouts observed at various field sites across India happened in absence of any females around the focal displaying male. Moreover, the presence of other males (adults and/or sub-adults) around the focal male increased the frequency of displays. The results indicate that displays were used not just for attracting females and/or as a signal for female mate choice but also potentially for establishing and keeping claim on display territory, to prevent other males from occupying that territory. Longer display bouts were recorded during May, June and July with the peak in the month of May. This phase of the breeding season (April-May-June) includes establishing and defending display territory from rival males. The weather is extremely dry and hot across different parts of India during May as monsoon rains arrive only in June or July. Although females do occasionally visit displays and display bouts reach their peak in May, mating rarely happens until after the first showers of Monsoon rains in June (DAP, personal observations). Thus, males spend most of their active time and energy during the month of May in displays often without food, water and mating. Some characteristic calls are also given at the beginning or end of the display and for advertising/ defending display territory (unpublished data). The display-related calls were also heard more frequently during May compared to the later part of the breeding season (VM and DAP, manuscript in preparation). As mentioned earlier, the displays in presence of females have certain additional elements (train rattle, bending forward with train up, hoot calls with mating attempts) while some features of the display (turns, wing-shake, display associated/ territory advertisement calls) are more frequently or exclusively seen in absence of females. Thus, one can argue that displays during the early part of the breeding season play a role in territory advertisement/ defence and male-male competition in this species.

Earlier studies from native and non-native habitats report that Indian peafowl form lek type of mating aggregations during the breeding season (Petrie et al., 1991; Yasmin and Yahya, 1996; Loyau et al., 2007). A typical lek consists of an aggregation of males in a specific arena where each male defends a small territory during the breeding season. Females have an opportunity to choose mates when they visit the arena. (Bradbury 1981). In a typical lek, the display sites or territory of males contains no significant resources required by the females except the males themselves (Bradbury 1981). The results of our study suggest that the choice of display sites which the males defend throughout the breeding season may not be random. In general, display sites were established at locations which had a small mound/ tree/ thicket or a place to perch when not displaying. About 86% of display sites had a medium to large-sized tree close by that provided protection from the sun and from potential predators such as feral dogs whereas, 28.67% of display locations were in a corner or in front of a stone wall/ mud wall/ some structure /tree that could provide a perch and also protection from wind when the display was going in slightly windy weather.

The display territories were not randomly or evenly distributed across the landscape. Seven distinct clusters of display territories were mapped at Morachi Chincholi over the period of study (Fig 1). However, these clusters do not conform to some of the criteria of a ‘classic lek’. Firstly, the area covered by clusters of display sites almost entirely overlapped with their normal movement range during the non-breeding season (Fig. 4). Thus, the clustering of display sites may be primarily due to the availability of patchy resources (food, water) rather than the aggregation of males strictly for breeding purposes. This situation does not conform to one of the defining characteristics of a classic lek. Although there were as many as 39 (and possibly more unmapped) display territories in some clusters, not all males in a cluster were within visible distance from each other. Topography, trees, shrubs, growing crops or simply distance between two display territories did not allow displaying males as well as visiting females to see all (or most) individuals at a time. Considering average distances between display territories and the area each cluster occupied (Table 2), these clusters can be potentially described as ‘exploded leks’ (Emlen and Oring 1977) or resource-based leks (Alexander 1975; Stiles and Wolf, 1979) since most of the display territories in Morachi Chincholi were close to food and/ or water sources. The movement of females near food provision sites and/ water resources may increase the chances of encounters between females and territory holding males (Fig. 4). The resource-based leks are more common in insects (Alexander 1975; Thornhill and Alcock, 1983) where breeding males form leks close to resources useful for females. This increases the chances of encountering females but males may not control access of females to these resources (Hoglund et al., 1998; Jiguet et al., 2002; Hingrat et al., 2008). The display territories that were occupied for more than one year during the study were more likely to be closer to a food or water source such as a farm, well, irrigation canal or typical female movement routes.

**Fig. 4:**
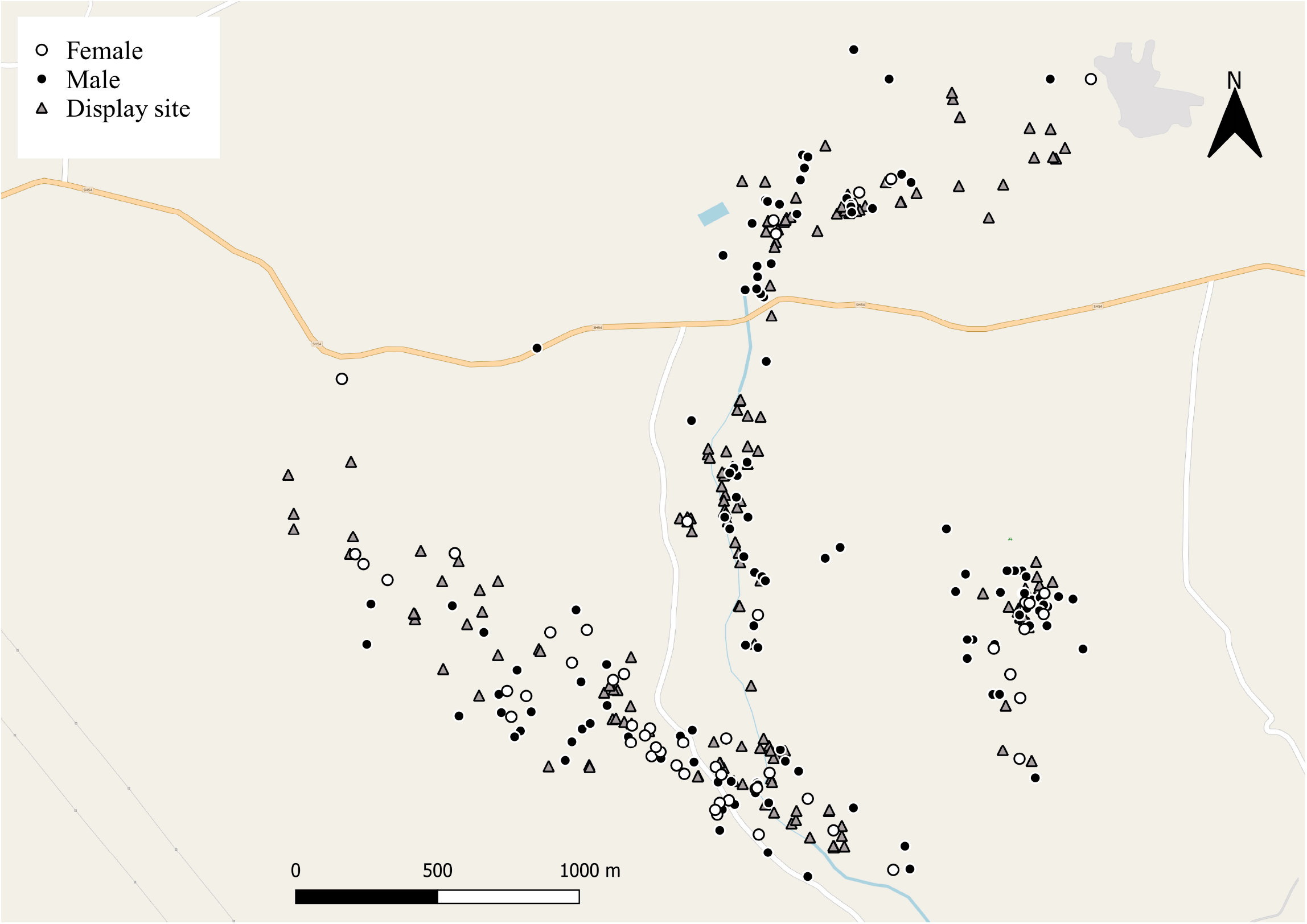
Distribution of display sites (grey triangles) in Morachi Chincholi completely overlaps with the area where males (black circles) and females (open circles) could be sighted during non-breeding season.

Establishing and holding a display territory at a prime spot (closer to resources such as food, and water with good protection from sun, wind and predators, closer to areas of female movement) requires the male to spend most of his active time during the breeding season at the display site often without food. At Morachi Chincholi, the display sites were concentrated within 300-500m around food provision sites (Table 3). The display sites were often established on the periphery of farms lining a small irrigation canal that runs through the village or close to other water sources such as well or percolation ponds. About 63.1% of display sites (118 out of 187) were within 500m of a water source (Fig 3). The vicinity of food and water sources gives the males easy access to food and water between display bouts without going far away from the display site (Fig. 3) as well as higher chances of encountering females who come for feeding (Fig 4). As males spend most of their active time during the breeding season at their display sites in vigilance, defence of display territory, maintenance and displaying (Harikrishnan et al., 2010 and personal observations), having a display site closer to food and water resources might be quite beneficial for the males. Therefore, this population might be considered to form exploded or resource-based leks.

Spatial distribution of leks close to food provision sites might also be beneficial for females as it reduces the cost of visiting leks and thereby the cost of making a choice as the females can assess potential mates while moving to feed (Loyau et al., 2007). However, to make the process of mate selection ‘economical’ the females should be able to assess the male displays almost simultaneously or sequentially within a short span of time which allows for quick comparison. The area covered by most clusters in our study was quite large and the distances between individual display sites were relatively greater for assessing the male displays almost simultaneously even if the females were moving through the area for foraging. Moreover, more than 3-4 males displaying simultaneously were rarely seen. Most likely, more than 4 males in a cluster did not display together but, we do not have sufficient data to support this due to difficulty in simultaneous tracking of several males in a cluster. However, they were within hearing distance of some of the other males. This might point to the exploded lek scenario. Neither male attractiveness nor the spatial distribution of display territories on exploded leks depends on resources or males establish display territories in the areas that maximise female encounters, in accordance with the hotspot model of lek evolution (Bradbury and Gibson, 1983; Jiguet et al., 2002). Male attractiveness was not measured in this study however, the spatial distribution of display territories was clearly influenced by resource availability (Table 3). As mentioned earlier, the vicinity of important resources during the breeding season might be beneficial for males in two ways-1. It potentially decreases the cost of searching for food while leaving the display territory unattended and 2. There are more chances of encountering females that are coming to the food/ water source. The data, therefore, points more towards a resource-based lek than a true exploded lek.

Although we are not sure whether the same individuals defended these sites across multiple years (as there was no individual identification/ tracking of males across years), previous studies (Loyau et al. 2007) show that peacocks show very high site fidelity across years with respect to establishing display sites. These sites were most likely the “hot spots” considering the vicinity of food and water resources and most importantly female movement. Similar observations have been noted by Loyau et al. (2007) in a feral population of Indian peafowl in France where non-defendable resources affect lek spatial organization in the peafowl. Females preferentially visited and mated with males displaying close to the food resources. Consequently, males competed to settle on those particular display sites where the probability to encounter females was higher (Loyau et al. 2007). The same study pointed out the need to investigate lek spatial organisation in wild populations of Indian peafowl as has been done in the present study. We could not conduct male removal experiments or measure the mating success of males closer (and farther) to food provision sites or water resources. The spatial distribution of display territories closer to food and/or water resources (Table 3, Fig 3) and some of the sites closer to resources being occupied in multiple years indicates that the peafowl population at Morachi Chincholi may form resource-based leks.

Simultaneous use of two mating strategies, i.e., exploded lekking and resource defence has been reported in a population of great bustards *Otis tarda* (Alonso et al., 2012). Resource-based leks have also been reported in bats *Mystacina tuberculata* (Toth et al., 2015) where males establish ‘singing roosts’ closer to communal roosts which female bats frequently use. In this species of bats, females are highly mobile and dispersed over a large area, therefore communal roosts dominated by females are considered a mobile resource and males try to establish leks closer to these communal roosts. In species mating in exploded leks, variability between males in the use of resource defence can create an intraspecific continuum between true lek and resource defence and such variability may occur in more species than realised earlier (Alonso et al., 2012). A striking difference between lekking ungulates and birds is that, in most lekking ungulates, there is considerable variation in mating strategies among and within populations. Males may hold resource-based territories and may also attempt to court and mate with females in mixed-sex herds (groups containing adults of both sexes (Höglund and Alatalo, 1995).

The increasing number of studies provide evidence that females choose their mates based on multiple cues rather than relying on single or few physical characteristics of males (Candolin, 2003). Multiple cues may provide additional information and may serve as multiple signals that indicate either general mate quality or enable females that differ in mate preferences to choose the most suitable mate (Candolin, 2003). Display sites may also play a role in male-male competition, in addition to the dynamic display (and calls) by the male who holds the territory throughout the breeding season as it allows greater chances of encountering females, similar to what is seen in a feral population of Indian peafowl in France (Loyau et al., 2007) or Lake Malawi chichlid fishes (Genner et al., 2008). Thus, display sites can be considered as a trait enhancing male competitive ability and may be favoured by females indirectly if it allows males to hold territories of the type and/or location visited more often by females (Wong & Candolin, 2005). Territories may indicate the availability of food and predation risk(s) faced by males, whereas male traits reflect the phenotypic and probably genetic quality of the partner (Balmford, Rosser & Albon, 1992). A correlation between male phenotypic traits and territory characteristics (or nest or other resource characteristics) has been found in several species, which implies that male and territory (resource) traits often back each other up as signals of male quality (e.g., Price, 1984; Kvarnemo, 1995; Bart & Earnst, 1999; Candolin & Voigt, 2001).

In summary, the current study points to the additional role of Indian peafowl male displays in territory advertisement and/or defence, especially during the early part of the breeding season. It also raises the possibility that Indian peafowl might form resources-based leks rather than classic leks as reported in earlier studies. Further studies for mapping the movement of individuals through breeding and non-breeding season as well as measuring female encounter rates and mating success of males closer (and farther) from resources such as food and water may strengthen the conclusions of this study regarding formation of resource-based leks.

## Significance statement

The role of extravagant display of Indian peafowl in female choice has been the focus of many previous studies in the native and non-native populations of peafowl. The results of this study on native peafowl populations suggest that the display and associated behaviours may have an additional role in male-male competition and/or territory defence during the breeding season. Also, the species may be forming resource-based leks rather than classic leks as thought earlier.

## Authors’ contribution statement

DAP conceptualized the study, acquired funding, was involved in data collection, and analysis and wrote the manuscript. VRM conceptualized study, collected field data, and was involved in data analysis and manuscript writing. PD collected field data.

## Data archiving statement

The data will be archived in the Dryad repository.

## Ethics statement

This study did not involve handling of any animals or collection of any type of samples from animals. The observation-based study followed all appropriate ethics protocols for fieldwork.

## Acknowledgements

Thanks to Dept. of Biodiversity, Abasaheb Garware College, Pune for hosting the fellowship and providing the infrastructural support. We are grateful to the villagers of Morachi Chincholi, Dr. Dharmendra Khandal, Meenu Dhakad and volunteers of Tiger Watch for hosting us and helping during the fieldwork. The authors are indebted to Prof. Milind Watve for the discussion which helped shape this manuscript better.

